# De novo linear phosphorylation site motifs for BCR-ABL kinase revealed by phospho-proteomics in yeast

**DOI:** 10.1101/2022.12.05.519126

**Authors:** Martin Smolnig, Sandra Fasching, Ulrich Stelzl

## Abstract

BCR-ABL is the oncogenic fusion product of tyrosine kinase ABL1 and a highly frequent driver of acute lymphocytic leukemia (ALL) and chronic myeloid leukemia (CML). The kinase activity of BCR-ABL is strongly elevated, however changes of substrate specificity in comparison to wild-type ABL1 kinase are less well characterized. Here, we heterologously expressed full-length BCR-ABL kinases in yeast. We exploited the proteome of living yeast as *in vivo* phospho-tyrosine substrate for assaying human kinases specificity. Phospho-proteomic analysis of ABL1 and BCR-ABL isoforms p190 and p210 yielded a high-confidence dataset of 1127 phospho-tyrosine sites on 821 yeast proteins. We used this data set to generate linear phosphorylation site motifs for ABL1 and the oncogenic ABL1 fusion proteins. The oncogenic kinases yielded a substantially different linear motif when compared to ABL1. Kinase set enrichment analysis with human pY-sites that have high linear motif scores well recalled BCR-ABL driven cancer cell lines from human phospho-proteome data sets.

**Highlights:**

- Full-length BCR-ABL kinase expression in yeast
- 1,127 pY-sites on 821 yeast proteins originating from ABL1 and BCR-ABL kinase activity
- Distinct linear kinase motif forABL1 and BCR-ABL p190 and p210 isoforms
- Validation of the kinase motifs through KSEA of human cancer cell line phospho-proteome data

**Significance:** The fusion protein BCR-ABL is a predominant oncogene in leukemia observed at very high frequency in chronic myeloid leukemia (BCR-ABL p210) and B-cell acute lymphoid (BCR-ABL p190) leukemia. In both cases the malignant transformation is reliant on elevated tyrosine kinase activity, however ABL1 kinase activation as such is not sufficient. At least in part, leukemias may therefore be driven by a specificity switch of the fusion kinase in comparison to the wild-type ABL1.

We investigated BCR-ABL kinase substrates in yeast through phospho-proteomics and established linear kinase phosphorylation-motifs for the full length BCR-ABL isoforms p210 and p190. These novel motifs differ substantially from the corresponding specificity determinants of wild-type ABL1 kinase, providing direct experimental evidence for altered phosphorylation-specificity of the oncogenic kinase fusion proteins.

## Introduction

The 90 human protein tyrosine-kinases (TKs) are key effectors of signal-transduction pathways. Tightly regulated, TKs mediate development, growth and multicellular communication in metazoans ^1^. However, perturbation of TKs results in deregulated kinase activities and malignant transformation of cells. Genetic alterations that can lead to activation of TKs involve gene amplification, missense variation, protein truncations and gene fusions through translocation of chromosomes. For example, chronic myeloid leukemia (CML) and a subset of acute lymphocytic leukemia (ALL) are causally linked to the expression of BCR-ABL, a gene fusion product that arises on the Philadelphia chromosome, a translocation between human chromosomes 22 and 9 ^2^. Different versions of the fusion protein are associated with the different forms of leukemia. The BCR-ABL fusion leads to activation of ABL kinase, which is a major, maybe sufficient, driver for disease development ^3^. Kinase inhibition with imatinib (Gleevec) has become the standard therapy for CML ^4^. Second and third generation inhibitor in combination with the allosteric inhibitor asciminib, constitute the newest line of therapy largely preventing recurrence through resistance mutations in BCR-ABL ^5^.

The activation of ABL and BCR-ABL has been studied in detail ^6^. BCR promotes dimerization or formation of larger oligomers of BCR-ABL critical for ABL kinase activation. The coiled-coil region of BCR and Y177 are important for the oncogenic properties. Elevated ABL kinase activity leads to activation of growth promoting signaling pathways including RAS/MAPK, PI3K, and JAK/STAT pathways, prerequisite for transformation of cells ^7^.

The oncogenic mechanism that drives cancer, however cannot be explained by elevated enzymatic activity of ABL alone ^6,7^. For example, the role of ABL myristoylation site is somewhat enigmatic. Though myristoylation has an inhibitory effect on kinase activity ^8^, and is absent in BCR-ABL, it was reported to be required for the transforming activity ^6,9^. Also, as mentioned, the coiled-coil region of the BCR part is important for the transforming activity. Importantly, increased ABL1 activity shows only modest oncogenic potential, by far less than expression of the BCR-ABL oncogene ^10,11^. This is also supported through cancer genome sequence projects ^12,13^. While amplification is a frequent event in cancers with EGFR kinase activation, gene copy number increase of ABL kinase as such is not observed in patient samples. Together this evidence supports the notion that a “simple” ABL1 kinase activity increase is not the sole cause for malignant transformation.

Depending on the actual chromosomal translocation, different BCR-ABL protein isoforms are generated with the most common BCR-ABL forms being BCR-ABL p210 and p190, respectively. The isoforms are differently associated with CML or ALL. The p210-BCR-ABL isoform is causal in 90% of CML cases, while p190-BCR-ABL occurs in 20-30% of B-cell acute lymphocytic leukemia ^14,15^. This isoform conundrum ^16^ was addressed by two proteomics studies which compared binding partners of p190-BCR-ABL and p210-BCR-ABL and indeed found substantial differences ^17,18^. There is additional evidence that the PH domain of the BCR part plays an important role modulating p210-BCR-ABL signaling networks ^19^. However, despite the distinct interaction patterns revealed, a viable hypothesis is that the kinase domain of p210 and p190, even though identical in the ABL part of the fusion protein, phosphorylates different subsets of substrates and thus drives the specific cancer phenotype. How phosphorylation specificity of BCR-ABL is altered in comparison to ABL, or in comparison of its protein isoforms, remains a largely open question.

The primary amino acid sequence surrounding a phosphorylation site, referred to as linear kinase phosphorylation motif, is the predominant kinase specificity determinant studied to date ^20–23^. Kinase motifs are typically obtained by assaying purified kinases, or the respective kinase domains, with synthetic peptide libraries or arrays *in vitro*. Utilizing experimentally derived motif data, a variety of computational approaches for scoring linear kinase motifs in the proteome to predict putative substrate sites have been developed ^20,24–27^. Linding et al. (2007) improved motif-based predictions by including contextual information, mainly protein-protein interaction (PPI) networks in a machine-learning approach. *In vivo*, contextual features such as protein levels, localization, adaptor/scaffold binding, aggregation phase and interaction network features may be equally important in establishing (or preventing) a substrate kinase relationship ^28–30^. Current methodologies to experimentally define kinase substrate relationships are either based on *in vitro* techniques using synthetic peptides / peptide arrays or are low throughput e.g. KESTREL ^31^ or engineered kinases allele technology by Shokat et al. ^32^. Recent approaches combining motif-centric *in vitro* kinase phosphorylation reactions with endogenous phospho-proteomes, boost p-site identifications and provide some information towards potential kinase substrates ^33^. However, no BCR-ABL linear motif signature has been reported.

We leverage the model organism *S. cerevisiae* to assay human tyrosine kinases thereby largely removing the complexity of human kinase signaling in cancer cells ^29,34^. To do so, we expressed full-length human protein tyrosine kinases at very low levels in yeast and subjected the yeast proteome to mass spectrometry based phospho-proteomics. In this approach the proteome of the growing yeast cell served as *in vivo* model substrate for an individual human kinase, which is acting in an intact, crowded, cellular environment. Most notably, the unicellular eukaryote lacks bona fide PTK signaling ^35–37^, thus each and every phospho-tyrosine site can be unambiguously attributed to one human kinase.

We previously probed the phosphorylation pattern of yeast proteins of strains expressing 16 of the 32 human non-RTKs and observed that they were very distinct. The phospho-Y-sites were then efficiently recorded after immuno-affinity purification (IAP) with mass spectrometry, resulting a large phosphoproteomics dataset of 26152 pY-peptides from 60 mass spectrometry samples. The data amounted to ∼ 1400 pY-sites on ∼ 900 yeast proteins representing ∼ 3700 kinase-substrate relationships^29^. The *de novo* motifs generated from our yeast data performed comparably or better than the reported motifs in the literature ^22^ when benchmarked against known kinase-substrate relationships ^29^. Here we assessed the specificity of p210 and p190-BCR-ABL kinase fusion proteins and their wild-type counterpart ABL1 using yeast as a model substrate. Our data suggest specificity differences that are relevant in human potentially leading to altered protein substrates for the oncogenic kinases.

## Results

### A yeast system to assay active full length BCR-ABL phosphorylation activity

Exogenous tyrosine phosphorylation in yeast can be toxic ^34^. Our assays critically employed very low expression of the full-length human tyrosine kinases that would otherwise severely impair yeast growth (**Figure 1**). To do so, kinases with (n) or without (c) a nuclear localization sequence, were expressed in yeast under control of a weak, copper (Cu^2+^) inducible promoter ^38,39^. Therefore, phosphorylation levels can be adjusted through the addition of Cu^2+^ to the growth media. To characterize clinical versions of oncogenic BCR-ABL kinase fusion proteins, we cloned p210- and p190-BCR-ABL in our yeast expression systems and probed the full yeast lysate with the anti pY-4G10 antibody. Indeed, both large, full-length, clinical versions of the fusion protein (**Figure 1**, p190 = 1530 aa; p210 = 2031 aa) showed immunoreactivity patterns on western blots of yeast whole cell lysate. Impaired cell growth was observed upon stronger induction with at 150μM copper and higher (**Figure 1**). Both, the phenotypes and the immuno-reactivity are directly related to human kinase activity. Functional BCR-ABL p210 was previously expressed in insect cells with the baculovirus expression system ^40^. However, purification of these large proteins remains a challenge, which can be overcome using our experimental yeast (“in vivo test tube”) approach ^34^ enabling to study phosphorylation activity of BCR-ABL.

**Figure 1:**
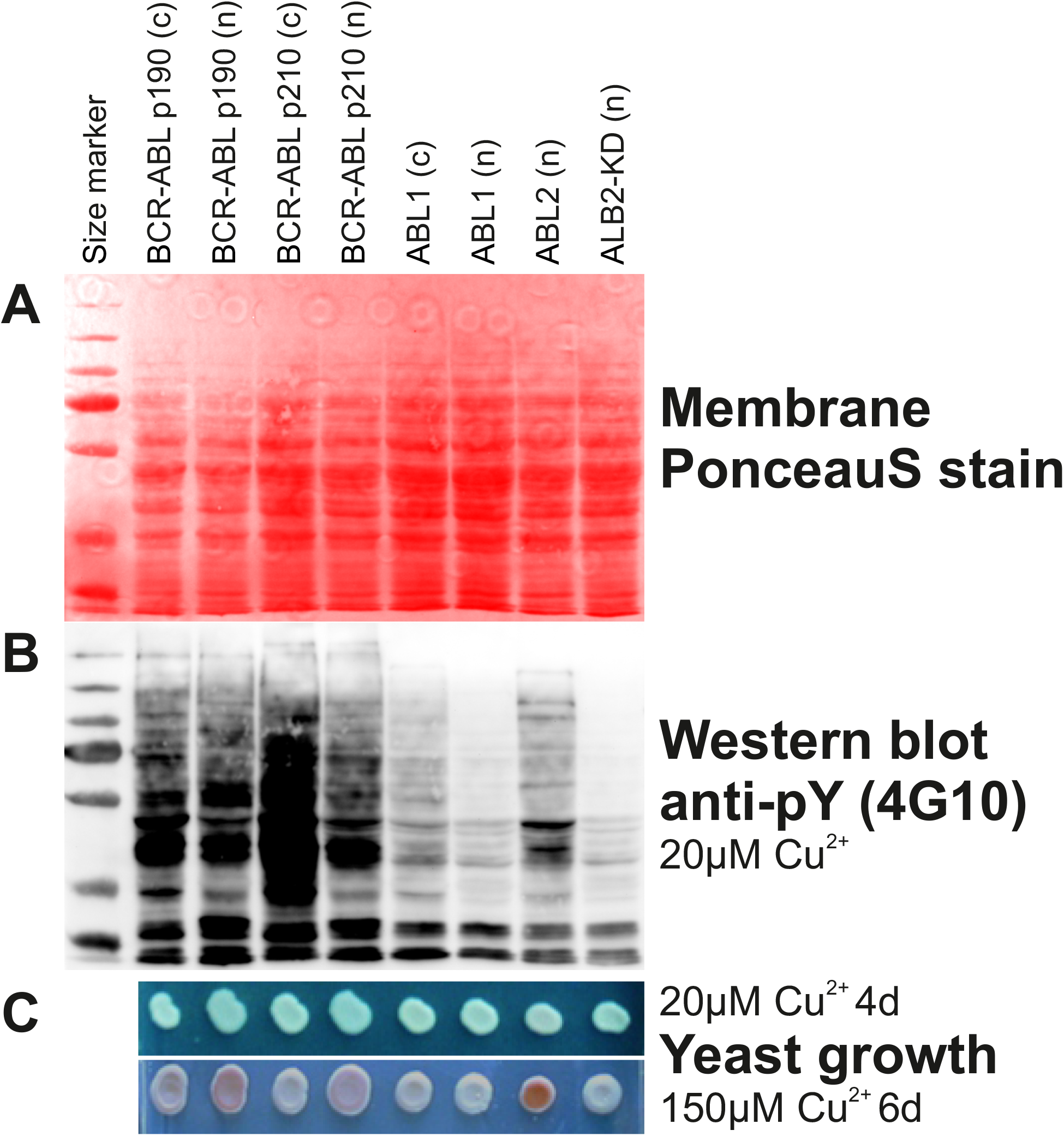
The large kinase fusion proteins BCR-ABL are active in yeast. ***A:*** Total cell lysates (kinases were induced with 20μM Cu^2+^) were stained on the membrane with PonceauS stain for loading control. ***B:*** Total cell lysates were probed with a pan anti-pY antibody (4G10, Sigma) and reveal human kinase activity in yeast. ***C*** *top:* Colony growth assay shows that low kinase induction (20μM Cu^2+^) does not affect yeast growth. ***C*** *bottom*: Induction with 150μM Cu^2+^ strongly affects yeast phenotypes of strains expressing active kinases. The red color phenotype is due to ade2 auxotrophy of the strains. [(n) = constructs with NLS, (c) = without NLS; ABL2 = pY positive control; ABL2-KD = kinase dead mutant version (K317M, P42684), negative control].

To obtain a large amount of input material for MS-based proteomics, yeast strains expressing ABL1 or BCR-ABL kinases respectively were grown in 5L cultures and prepared for LC-MS/MS. Because of the different levels of activity of the ABL1 wild-type and BCR-ABL kinases (**Figure 1b**) we prepared three cultures of ABL1(c), two of ABL1(n) and one each of the four BCR-ABL versions. In order to identify yeast proteins that were phosphorylated by human ABL1, p210-BCR-ABL and p190-BCR-ABL, respectively, we subjected 98% of the cell lysates to phospho-Y enrichment and 2% for input proteome analyses (**Supplemental Table 1**). The input samples demonstrated the expression of the individual human kinases with good peptide coverage (**Figure 2, Supplemental Table 2**). A large number of unique peptides corresponding to the human kinases was identified. The coverage of peptides in the corresponding protein regions including a p190 BCR-ABL breakpoint peptide and a series of isoform specific peptides confirmed expression of the full-length proteins in yeast.

**Figure 2:**
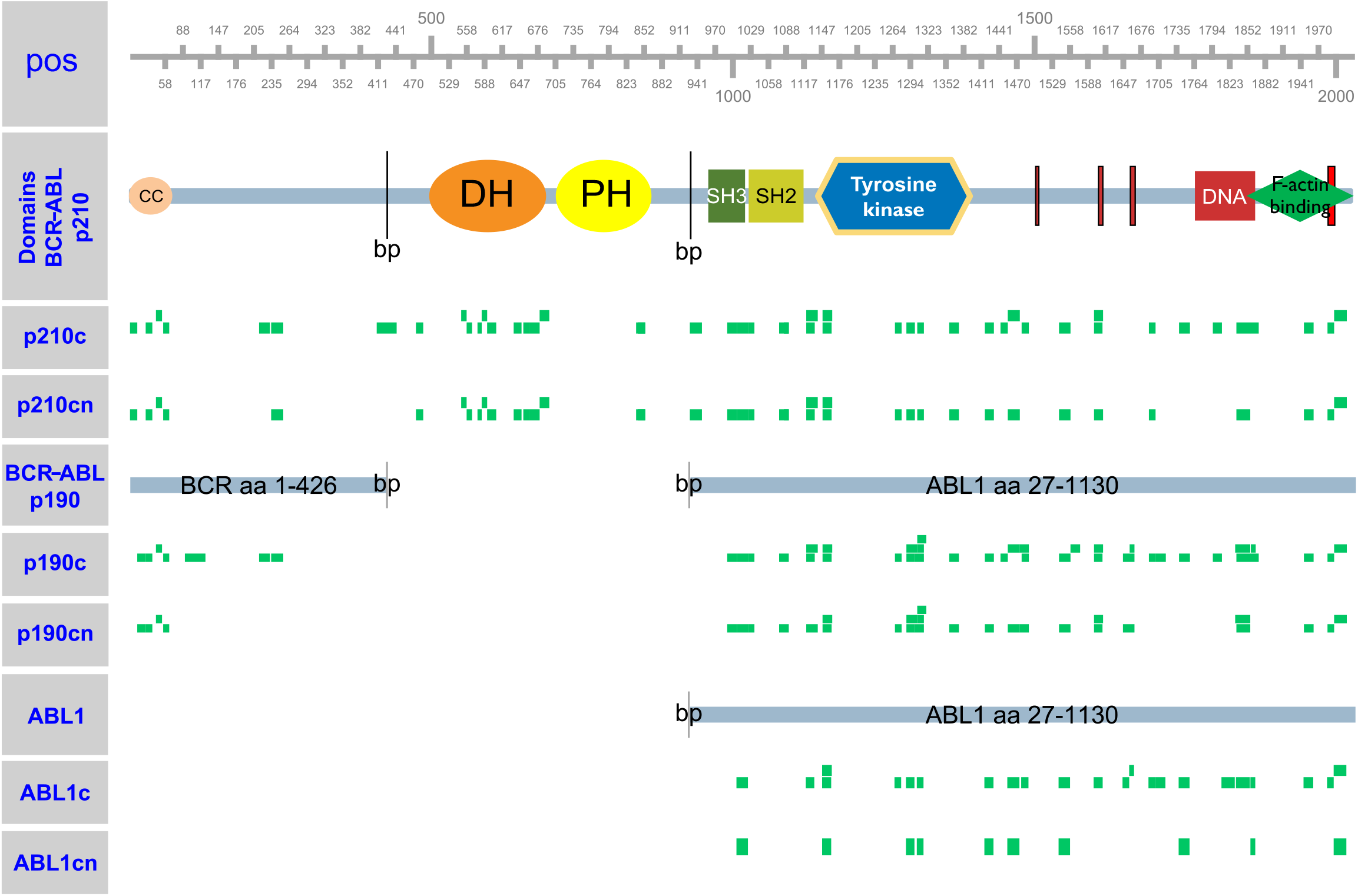
Peptide coverage of ABL1 and BCR-ABL kinases. Green rectangles, positioned according to the BCR-ABL or ABL1 sequence, indicate peptides identified from the yeast protein extract for a given strain. Notably, the p190 BCR-ABL breakpoint covering peptide (TGQIWPNDGEGAFHGDA|*EALQR*) and several p210 BCR-ABL specific peptides, such as the one covering the p210 breakpoint region (TGQIWPNDGEGAFHGDA|*DGSFGTPPGYGCAADR*) were detected.

The digested peptide samples were subjected to antibody based phospho-Y enrichment, mass spectrometry analyses, sequence identification (MaxQuant), and data processing (quality control and abundance filtering). To obtain a high-confidence data set, known endogenous yeast pY-sites ^37^ were removed. Secondly, pY-sites were compared to the Corwin dataset ^29^ where phospho-Y-proteomes for 16 human non receptor tyrosine kinases were recorded in a similar approach. Sites that also occurred in more than six preparations with different tyrosine kinases in the Corwin dataset were removed. Third, sites that were measured with more than six non-receptor tyrosine kinases in the Corwin dataset and were identified in 12 or more runs in this study were removed as unspecific. With this filter we removed a total 157 peptides corresponding to 1434 SPCs from the raw pY-peptide data. The final 1186 peptides covered 1127 unique pY-sites on 821 proteins and form our high-confidence pY-dataset, 611, 477 and 655 pY-sites were identified for ABL1, p190-BCR-ABL and p210-BCR-ABL respectively (**Figure 3a, Supplemental Table 3**).

**Figure 3:**
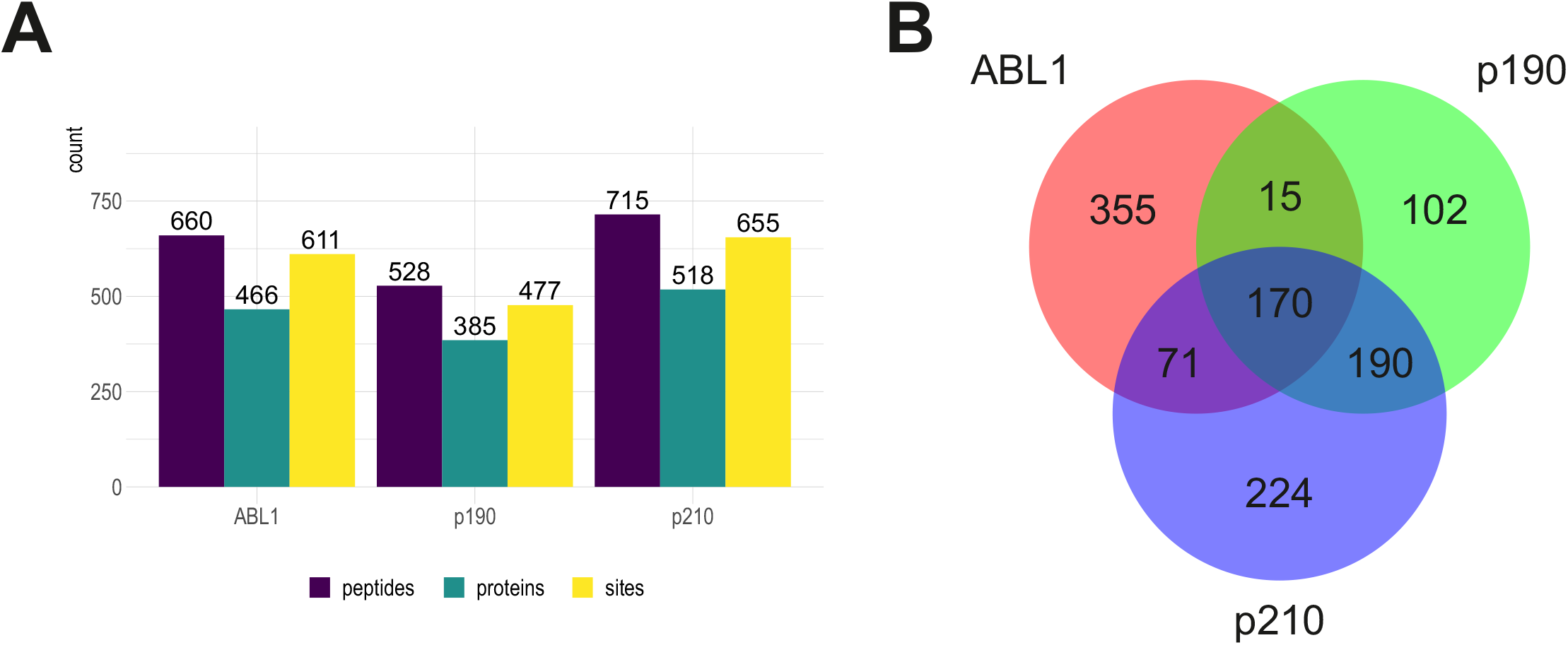
High confidence pY-dataset. ***A:*** pY-peptides, annotated proteins and pY-sites for ABL1, BCR-ABL p190 and BCR-ABL p210. ***B:*** Venn diagram showing the 1127 pY-sites overlap between ABL1, BCR-ABL p190 and BCR-ABL p210.

When comparing the pY-sites, we found that the overlap of BCR-ABL isoforms is larger than the overlap of each isoform with the wild-type ABL1 (Jaccard-similarity: ABL1 vs p190: 0.205, ABL1 vs p210: 0.235, p190 vs p210: 0.466). This suggests that in our yeast model system different protein tyrosine sites are phosphorylated by the different ABL1 kinases, and that the difference between wild-type ABL1 and the oncogenic BCR-ABL is most pronounced (**Figure 3b**).

### De novo kinase motifs for BCR-ABL

These data were then used to derive *de novo* linear sequence recognition motifs for ABL1 and BCR-ABL kinases. We applied iceLogo ^41^ to determine enriched amino acids at positions -7 to 7 surrounding the pY-residues (7Y7 mers). The background set for the analysis was generated from non-modified peptides with a tyrosine residue in the input samples (ABL1: 7857; p190: 7892; p210: 7864 sites). The ABL1 motif obtained agrees well with what was reported in the literature (**Figure 4A**) ^42^. The ABL1 motif is largely determined by a proline at the +3 position (P+3).

**Figure 4:**
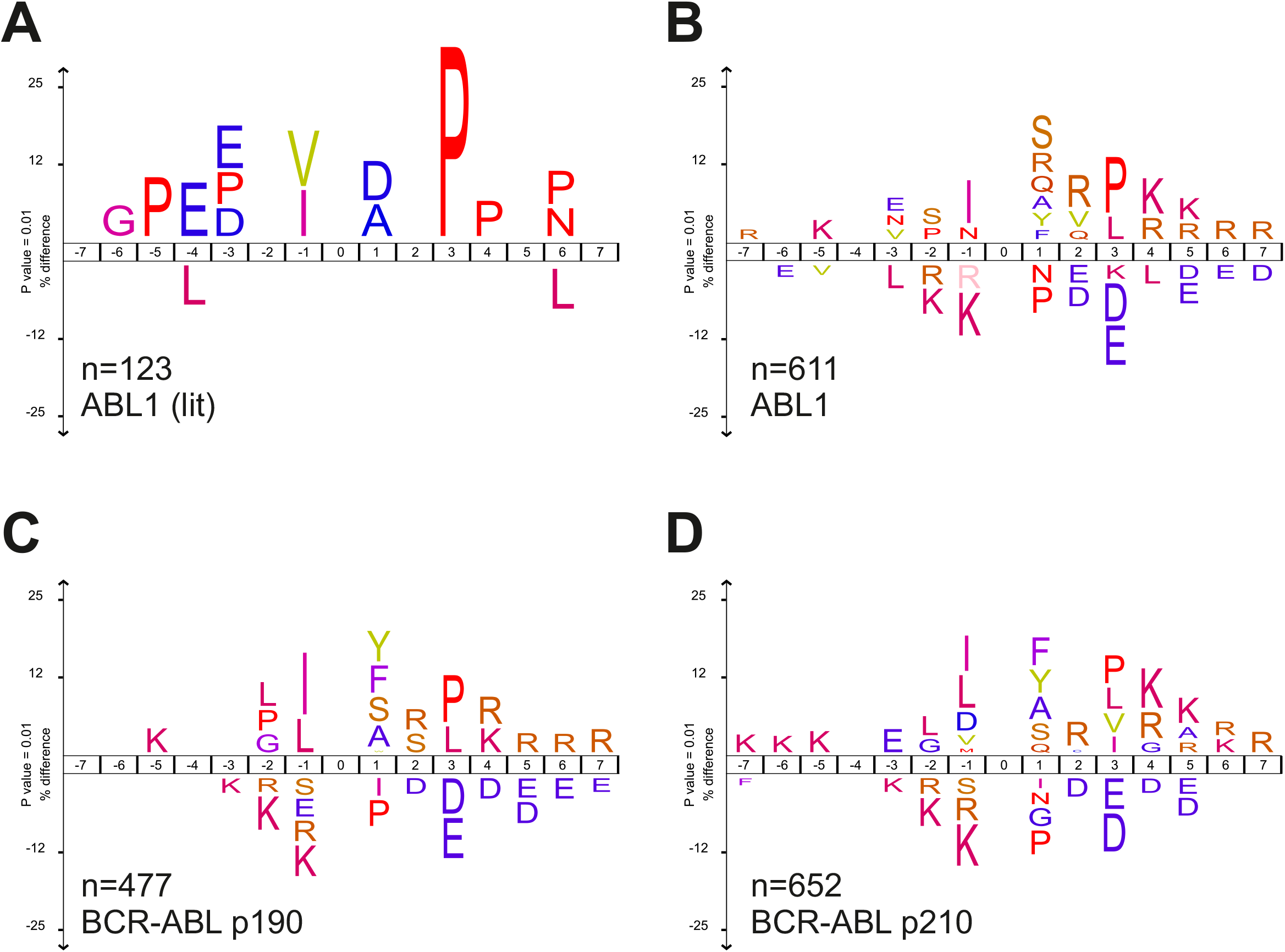
De novo kinase phosphorylation motifs of ABL1 and BCR-ABL isoforms. Kinase motifs were computed with iceLogo ^41^ (https://iomics.ugent.be/icelogoserver/). Number of input 7Y7-sequences (n) are indicated. ***A:*** ABL1 (lit): redraw of ABL1 literature motif ^42^. ***B-D:*** ABL1, p190 BCR-ABL and p210 BCR-ABL motifs derived from the yeast pY-data.

In agreement, the analysis of yeast de novo peptide-derived motifs of ABL1 (**Figure 4B**) similarly yields the highest value of 2.9392 for P(+3) in the position-specific scoring matrices (PSSM) (**Supplemental Table 4**). Interestingly, in both BCR-ABL isoforms, the P(+3) scores high (2.6597 in p190, and 2.0341 in p210), however it is no longer the main determinant with the highest score (rank 3/300 in p190 and rank 10/300 in p210). In p190, I(-1) with 3.4928 is followed by tyrosine Y(+1) with 3.3090 (**Figure 4C**). In p210, the position with the highest PSSM value is Y(+1) with 2.7834 followed by phenylalanine F(+1) with 2.6854 (**Figure 4D**). The data suggest that in the motif sequences of BCR-ABL in comparison to wild-type ABL1 determinants, the P(+3) has less weight and that the (-1) and (+1) position next to the Y-site contribute more strongly to phosphorylation specificity. While this finding requires further investigation, we note that to our knowledge this data represent the first specific linear sequence motif for the BCR-ABL oncogenic kinases.

### Scoring human phospho-proteome data with the BCR-ABL specific kinase motif

In order to test whether the generated motif reflects *in vivo* phosphorylation activity in human cells, we collected pY-proteomics data from the literature. Rikova et.al. ^43^ reported pY-proteomics of NSCLC cell lines and patient samples, including 423 pY-sites of the NSCLC cell line HCC78. Beekhof et al. ^27^ reported pY-proteomes of several cell lines, PDX and patient samples. This included 3911 pY-sites of a HCC827-NSCLC, as well as 1543 pY-sites of a K562 CML cancer cell line and 1724 pY-sites of a PDX-pY-IP colorectal cancer patient-derived xenograft. As a validation, we used the PSSM from our motifs to quantitatively score the pY-human datasets. Using the top scoring sites we queried the ranked phospho-tyrosine proteomes in a KSEA gene set enrichment analysis (adapting the code of ^44^). Using our de novo motif KSEA analysis for ABL1, the p190 and p210 BCR-ABL, we found that HCC78 phospho-proteome was enriched for ABL1 activity (**Figure 5**). HCC78 is a NSCLC cell line driven by a SLC34A2/ROS1 fusion and known to exhibit various elevated non-RTK activities including ABL1 ^43^. The K562-CML cancer cell line is driven by p210 BCR-ABL ^45^ and therefore KSEA enrichment indicates elevated ABL1 activity when probed with ABL literature substrates (not shown). Using our de novo motifs, strong enrichment was found for pY-sites that score very high with p210-BCR-ABL in contrast to ABL1 and p190-BCR-ABL. Results for the EGFR-driven cell line HCC827 and the KRAS-driven patient-derived colorectal cancer xenograft PDX-pY-IP are presented with negative results for comparison (**Figure 5**). The analysis showed that our motifs derived from BCR-ABL activity in yeast matches specific activities that can be profiled in phospho-proteomics measurement of human cancer cells.

**Figure 5:**
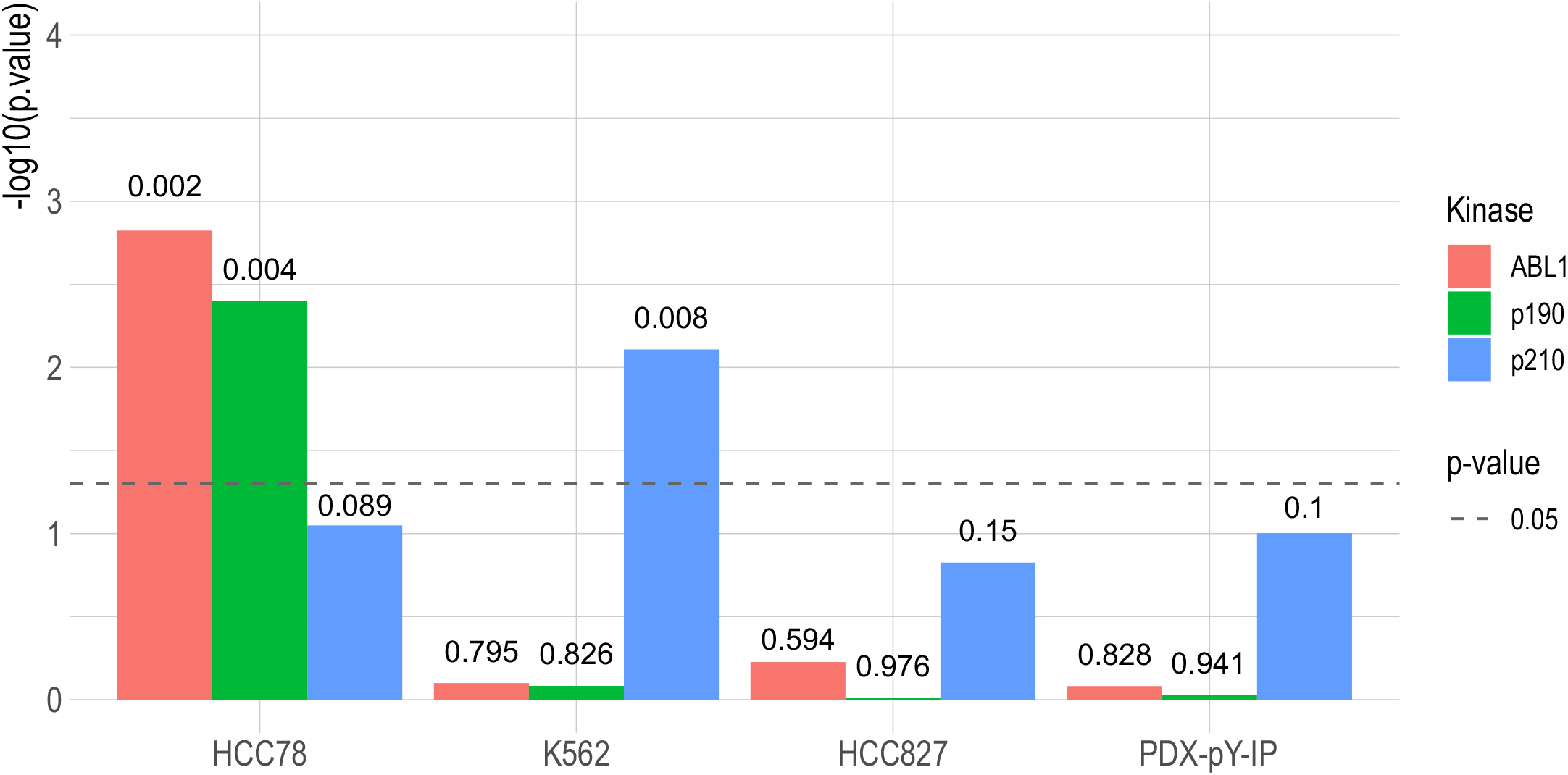
Motif-based KSEA results for human literature phospho-proteomics data. X-axis denotes biological source material (cell line or xenograft) HCC78: lung cancer cell line (n=423 sites); K562: BCR-ABL driven CML cell line (n=1543 sites); HCC827: NSCLC cell line (n=3911); PDX-pY-IP: colorectal cancer patient-derived xenograft (n=1724). Y-axis shows -log10-transformed p-value. Bar colors represent kinase (ABL1: WT ABL1, p190: BCR-ABL p190, p210: BCR-ABL p210), dashed line denotes p-value significance cutoff, numbers above bars show p-value. Query set was top 5% scored with the linear motifs of the respective kinases.

## Discussion

Major obstacles in defining kinase-substrate relationships stem from the fact that, at any point in time, kinases are differentially expressed dependent on cell type or cell-cycle phase or subcellular localization, exhibit partly overlapping substrate specificity, and have magnitude differences in enzymatic activity. Furthermore, kinases form complex signaling networks including redundancy and feedback loops. Together this hinders identification of kinase targets using classical perturbation approaches in the context of endogenous cellular signaling. The most instructive (because the most comprehensive) example to illustrate this is a study by Bodenmiller et al., who recorded high quality phospho-proteomes from 97 individual kinase yeast knock out strains ^46^. Inactivation of most kinases affected large parts of the phospho-proteome, not only the immediate downstream targets of the kinases. Roughly the same number of sites with decreasing phosphorylation (expected) and increasing phosphorylation (not expected) were found in the majority of knock out strains, so that essentially no kinase-substrate relationships could be truthfully established. In a similar, more recent study that again quantified phospho-proteomes of 80 kinase knockout strains in yeast 3724 phospho-sites decreased and 3501 phospho-site increases were revealed ^47^. Though these studies are examining yeast, the same observations hold for the complex phospho-signaling systems in mammalian systems.

When comparing the phospho-proteome of p190-BCR-ABL or p210-BCR-ABL cell lines with their parent counterparts, a total of 812 phospho-sites were quantified of which 302 stood out due to increased intensity of phosphorylation ^18^. Intriguingly the same number of phospho-sites showed decreased phosphorylation in the transformed cells, with the remaining ∼ 200 being unaffected. In a parallel proteomics study ^17^ roughly 1/3 of the pY-sites recorded decreased and 2/3 increased in the presence of BCR-ABL. Differences in phospho-signaling profiles of p190-BCR-ABL and p210-BCR-ABL in Ba/F3 and HPC-LSK cell lines, again pY-site decreases and increases, were also observed using a tyrosine phosphorylation antibody array covering 228 phospho-tyrosine sites ^48^. This illustrates that it is difficult to define direct kinase-substrate relationships through proteomics screens in mammalian cells. Complex signaling networks, feedback loops, conditional activity and kinase redundancy combine to confound systematic attempts to address direct kinase-substrate identification.

Here, we heterologously expressed the full-length human oncogenic BCR-ABL kinases, about 1500 and 2000 amino acids in size, in yeast exploiting the yeast proteome as an *in vivo* model substrate for tyrosine phosphorylation. In contrast to *in vitro* peptide array approaches ^21,22^, full length kinases were used and the substrate yeast proteome resembled a fully folded, crowded, competitive, cellular context. Very low kinase activity is required to avoid yeast growth defects ^34^, however yeast can be cultured in large volumes to obtain large amount of material for p-Y enrichment and mass spectrometry analysis. Therefore, we assigned 611, 477 and 655 pY-sites unambiguously to ABL1, p190-BCR-ABL and p210-BCR-ABL activities, respectively.

With this large data set we determined consensus phosphorylation site motifs targeted by ABL1 and BCR-ABL1. We note that only a minor fraction of the identified sites actually results any match when scored with the consensus motif at least partially. In agreement with this, 80% of the almost 300,000 human phospho-sites mapped today (https://www.phosphosite.org/staticSiteStatistics) do not match any known kinase motif, and conversely biochemical and bioinformatic approaches have only identified kinases for less than 5% of the phospho-sites in the human proteome ^49^. The predictive value of linear kinase motifs in general remains somehow limited, however no BCR-ABL linear motif signature has been reported.

Our motif results were validated in two ways. First, the ABL1 motif generated de novo with wild-type ABL1 very well resembles linear phosphorylation motif from the literature that was established on an individual basis through inspection of known ABL1 phosphorylation sites. Interestingly the motif generated from a similar number of phospho-sites was substantially different for the BCR-ABL fusion proteins when compared to wild-type ABL1. We observed less contribution to the specificity signature from the +3 proline and relative higher weights for scoring other positions at +1 and -1. This observation supports the hypothesis that the oncogenic versions of ABL1 does not only have higher activity but also encounters substrates with a different substrate specificity even though the kinase domain is identical in the proteins. This change of specificity could lead to aberrant phosphorylation of proteins and contribute to cancer development or progression in CML. We validate the motifs by KSEA of human phospho-proteome data from a K562 CML cell line, that is driven by expression of p210 BCR-ABL. As a result, we obtained significant enrichment of high scoring pY-phospho substrates with the corresponding p210 motif but not with the p190 or the ABL1 wild-type motif.

Our approach does not provide direct human kinase-substrate relationships, however using the living yeast cell as an *in vivo* substrate space has proven useful to address human kinase substrate specificity. Linear phosphorylation site motifs for kinases are key in kinase substrate predictive analyses ^50^, and we report such motifs for BCR-ABL. Moreover, our experiments provide first direct experimental data that demonstrate altered substrate specificity of the oncogenic BCR-fusion kinase in comparison to wild-type ABL1.

## Material and Methods

### Yeast strain, plasmids and transformation

The *Saccharomyces cerevisiae* strain L40ccU2 (U2) ^51^ was used for all experiments. Strains were transformed by the lithium acetate method with plasmids carrying the kinase of interest under a copper-inducible promoter. The pASZ-DM plasmids used here have been described previously ^38^.

### Yeast culture and harvest

Yeast was cultured in three steps to reach the main culture volume of 5L. First, a 10ml ONC of selective media (NB, 2% glucose, minus adenine) was inoculated (ONC1) and grown ON (20-24h) @30°C, 180rpm. Second, ONC1 was used to inoculate a 135ml culture (ONC2) that was grown ON again. Third, ONC2 was used to inoculate the 5L main culture to OD_600_=0.05. The main culture was then grown for 6h @30°C, 180rpm before addition of CuSO_4_ to a final concentration of 20μM (BCR-ABL) or 300μM (ABL1). After 20h cells were harvested by centrifugation (4°C, 4000rpm, 10min), aliquoted into portions of 1.5ml in lysis tubes, snap frozen in liquid nitrogen and stored @-80°C until cell lysis and further processing.

### Cell lysis, reduction, alkylation and tryptic digest

Cell pellets were mechanically lysed with zirconia beads in high-molar urea buffer (9M urea, 100mM AMBIC) in a volume ratio of 2+1+1 (cells, beads, buffer). Lysis was performed in a SPEX Geno/Grinder® (3x 2min agitation at 1500rpm with 15s pauses). Samples were centrifuged (4°C, 15000rpm, 15min) and the supernatant was moved to a fresh tube. Lysis was repeated twice by adding fresh buffer. Cleared lysates were combined and subjected to reduction (TCEP, 5mM final concentration, 60°C, 20min) and alkylation (chloroacetamide, 10mM final concentration, room temperature, 10min in the dark). To reduce urea concentration, samples were diluted 4.5x with 100mM AMBIC before tryptic digest (in-solution, overnight at room temperature, protease/protein ratio 1/100 w/w).

### C18 column purification

Peptide lysates were acidified (trifluoroacetic acid (TFA), final concentration 1%, 10min at room temperature), centrifuged (4000rpm, 5min, @RT) and purified by C18 vacuum column (Waters). The column was pre-wet with 5ml 100% acetonitrile (ACN) and washed twice with 3.5ml 0.1% TFA (solvent A) before lysate loading. The entire sample was processed in one session, the column was then washed consecutively with increasing volumes of solvent A (1ml, 5ml, 6ml) before peptide elution. Peptides were eluted using 3x2ml of solvent B (0.1%TFA + 40% ACN). Each sample was split into two aliquots, one with 98% volume for immuno affinity enrichment of phospho-peptides and one with 2% volume for total peptide analysis. Both aliquots were snap frozen in liquid nitrogen for at least 30min and subjected to lyophilization before further processing.

### Enrichment of pY-tryptic peptides

Peptides were then subjected to pY-peptide enrichment. For anti-phospho-tyrosine immuno-affinity enrichment we built on the protocol first established by Rush et al. ^52^. The P-Tyr-100 PhosphoScan Kit was obtained commercially (Cell Signaling Technology, Danvers, MA, USA). The procedure was performed by manufacturer’s instructions including peptide purification with C18 ZipTips® (Merck). The eluted peptides were snap frozen and lyophilized before reconstitution in 0.1% formic acid for HPLC-MS/MS.

### Mass spectrometry

LC-MS/MS analyses were performed on a TimsTOF Pro (Bruker) equipped with an UltiMate™ 3000 RSLCnano System (Thermo Fisher Scientific) run with an Aurora C18 column (Ion Opticks). The mass spectrometer was run in positive ion polarity mode, data-dependent acquisition (DDA), TIMS and PASEF. Instrument control and data acquisition was handled by otofControl v3.6. LC/MS workflow automation was handled by Bruker Compass HyStar 5.1. Data post processing was handled by DataAnalysis software.

Mass Spec raw files (.d) were analyzed with the quantitative proteomics software MaxQuant (v1.6.15.0). MaxQuant was run with default parameters with the following adaptations. In the section “group-specific parameters” the option “label-free quantification” (LFQ) was enabled. Trypsin was selected as the sole protease and up to two missed cleavages were allowed. Cysteine carbamidomethylation (+57.021464 Da) was assigned a fixed modification due to the alkylation step. Variable modifications were methionine oxidation (+15.994915 Da), N-terminal acetylation (+42.010565 Da) and tyrosine phosphorylation (+79.966330 Da). FDR was set to 1% and decoy peptides were generated by reversing the theoretical spectra from the FASTA file. Minimal peptide length was set to 7. The reference FASTA file was the UniProt complete *Saccharomyces cerevisiae* proteome (reviewed, release October 2020, 6721 entries) with select human FASTA sequences added for human ABL1, BCR-ABL p190 and BCR-ABL p210.

### Computational analyses

Analysis of MaxQuant output was performed using bioinformatics program packages including R (Statistics, enrichment analyses) and custom-made tools in Perl or SQL. Kinase consensus motif generation was performed using iceLogo ^41^. Motif scores were calculated in R based on the iceLogo position-specific scoring matrices (PSSM). Kinase substrate enrichment analysis (KSEA) was performed in R using the code of David Ochoa ^44^.

### Motif generation

Motif scores were generated from the amino-acid frequency PSSMs generated by iceLogo. First, the amino acid frequency ratio was determined for each amino acid at each position in the 7Y7 motif. This yields a 20x15 matrix for each kinase (20 AAs, 15 positions). Second, the motif score for any 7Y7-motif is calculated by summing up the amino acid frequency ratio of the respective amino acids at the respective position in the motif. This yields one score for each motif per kinase.

### Kinase substrate enrichment analysis (KSEA)

To validate our motif score we tested it against literature phospho-proteomic datasets ^27,43^ using the KSEA algorithm ^44^. First, the literature datasets are ranked by the provided MS metric (intensity or spectral counts). Second, every pY-peptide from the literature dataset is converted to a 7Y7-motif. Third, each motif is assigned a score based on the linear motif of the kinase (ABL1, p190 or p210). Fourth, the top 5% by motif-score are defined the query set for KSEA.

KSEA calculates enrichment of a subset of items (query set) in a larger reference set. In our setup all the elements of the query set are part of the reference set since we select a subset. The ranks of the motifs in the query set within the peptides of the reference set are determined and a running sum-based enrichment score (ES) is calculated. Statistical significance is determined by comparing the experimental ES to an ES-list generated from randomized null distributions. The resulting ES and p-value inform on enrichment or depletion of the query set within the reference set and its statistical significance.

## Data accessibility

The mass spectrometry proteomics data have been deposited to the ProteomeXchange Consortium via the PRIDE partner repository with the dataset identifier PXD038551.

## Acknowledgements

We thank Evelyne Jany-Luig for help with the cloning of BCR-ABL constructs. We thank Thomas G. Graeber (UCLA) for providing the BCR-ABL cDNAs. The work was supported by the Austrian Science Fund through FWF project P30162 and through the FWF doc.fund Molecular Metabolism (DOC 50). MS proteomics was supported by the Field of Excellence BioHealth—University of Graz.

## Author contributions

Martin Smolnig: Methodology, Software, Validation, Formal analysis, Investigation, Data curation, Writing – Original Draft, Writing – Review & Editing, Visualization, Project administration.

Sandra Fasching: Investigation.

Ulrich Stelzl: Conceptualization, Formal analysis, Resources, Writing – Original Draft, Writing – Review & Editing, Supervision, Funding acquisition.

